# ANGPTL8 R59W variant influences inflammation through modulating NF-κB pathway under TNFα stimulation

**DOI:** 10.1101/2023.07.04.547624

**Authors:** Mohamed Abu-Farha, Dhanya Madhu, Prashantha Hebbar, Anwar Mohammad, Arshad Channanath, Sina Kavalakatt, Nada Alam-Eldin, Fatima Alterki, Ibrahem Taher, Osama Alsmadi, Mohammad Shehab, Hossein Arefanian, Rasheed Ahmad, Fahd Al-Mulla, Thangavel Alphonse Thanaraj, Jehad Abubaker

**Author notes:** **Correspondence:** Jehad Abubaker, PhD, Department of Biochemistry & Molecular Biology, Dasman Diabetes Institute, P.O. Box 1180, Dasman 15462, Kuwait, Thangavel Alphonse Thanaraj, PhD, Department of Genetics and Bioinformatics, Dasman Diabetes Institute, P.O. Box 1180, Dasman 15462, Kuwait, & Fahd Al-Mulla, MD, PhD, Department of Genetics and Bioinformatics, Dasman Diabetes Institute, P.O. Box 1180, Dasman 15462, Kuwait. M.A-F and DM. contributed equally to this work.

## Abstract

**Background:** ANGPTL8 is known to regulate lipid metabolism and inflammation. It interacts with ANGPTL3 and ANGPTL4 to regulate LPL activity, and with IKKα/β to modulate NF-κB activity. Further, a SNP leading to ANGPTL8 R59W variant associates with reduced LDL/HDL and increased FBG in Hispanic and Arab individuals, respectively. In this study, we investigate the impact of R59W variant on the inflammatory activity of ANGPTL8.

**Methods:** ANGPTL8 R59W variant was genotyped in a discovery cohort of 867 Arab individuals from Kuwait. Plasma levels of ANGPTL8 and inflammatory markers were measured and tested for associations with the genotype; the associations were tested for replication in an independent cohort of 278 Arab individuals. Impact of the ANGPTL8 R59W variant on NF-κB activity was examined using approaches including overexpression, luciferase assay, and structural modeling of binding dynamics.

**Results:** The ANGPTL8 R59W variant was associated with increased circulatory levels of TNFα and IL7. NF-κB activity, as assessed by the increased in the phosphorylation of IKK-α/β protein, IκBα, and NF-κB p-65 in R59W variant compared to wild type, and TNFα stimulation further elevated it. This finding was substantiated by increased luciferase activity of NF-κB p65 with the R59W variant. Modeled structural and binding variation due to R59W change in ANGPTL8 agreed with the observed increase in NF-κB activity.

**Conclusion:** ANGPTL8 R59W is associated with increased circulatory TNFα, IL7 and NF-κB p65 activity. Weak transient binding of ANGPTL8 R59W variant explains its regulatory role on the NF-κB pathway and inflammation.

## 1 Introduction

Obesity and Type 2 Diabetes (T2D) have become serious global health conditions that have reached alarming levels ^1^. They are associated with impaired metabolism leading to high circulating free fatty acids (FFAs) and triglycerides (TG) ^2^. Another outcome of T2D is the inability of pancreas to meet the increased insulin demand leading to high glucose concentration in the blood circulation ^3^. A family of proteins that is structurally similar to angiopoietins has been identified as angiopoietin-like proteins (ANGPTLs) and it has been demonstrated to play an important role in gluco-lipid homoeostasis. They are a group of secreted glycoproteins composed of eight members (ANGPTL1-8). All of them contain an amino-terminal, linker region and a carboxy-terminal fibrinogen-like domain. However, ANGPTL8 lacks fibrinogen/angiopoietin-like domain ^4^. ANGPTL8, also known as Lipasin, RIFL, TD26, or Betatrophin, is a 22 KDa protein comprising 198 amino acids and is a distinct member of angiopoietin-like proteins family. Many of the ANGPTLs are involved in multiple biological processes such as lipid metabolism, inflammation^5^, and hematopoietic stem cell activity ^6,7^.

ANGPTL8 is abundant in liver and fat tissues particularly in brown adipose tissue (BAT) and white adipose tissue (WAT) ^8^. Nutritional regulation has been advocated as an important component on ANGPTL8 expression ^9^. During fasting, ANGPTL8 expression was reduced by 80% in BAT and WAT whereas mice exposed to high fat diet significantly increased ANGPTL8 mRNA expression in liver and BAT ^8^. In human obese subjects with metabolic syndrome, plasma ANGPTL8 levels have been inversely correlated with protein intake ^10^. Hence ANGPTL8 level is greatly modulated by the amount of protein intake. During fasting conditions, glucocorticoids are increased; and they suppress ANGPTL8 expression through activation of glucocorticoid receptor that binds to the negative glucocorticoid response element (nGRE) ^11^.

Several studies have reported that ANGPTL8 acts as a key modulator in lipid metabolism and metabolic disorders ^12,13^. ANGPTL8 has at least two physiological roles -namely those of regulating plasma TG levels and controlling inflammation. Overexpression of ANGPTL8 in mice increased TG levels more than five folds ^8^ whereas its deficiency reduced TG by two folds ^14^. ANGPTL8 deficiency increases postprandial lipoprotein lipase (LPL), especially in cardiac and skeletal muscles, and a proposed mechanism was shown to demonstrate ANGPTL8 on TG metabolism in different nutritional states wherein ANGPTL3, ANGPTL4 and ANGPTL8 coordinate to regulate TG trafficking ^15^. Research has shown that circulating levels of ANGPTL8 were elevated in arteriosclerosis and non-alcoholic steatosis ^16,17^. Plasma levels of ANGPTL8 were seen elevated in patients with severe infections. Strong correlations between circulating ANGPTL8 and LPS-induced acute inflammatory response have been demonstrated in animal models ^18^. Henceforth ANGPTL8 and its impact on regulation of blood lipoproteins throws light on its role in modulating inflammation.

Nuclear factor kappa-light-chain-enhancer of activated B cells (NF-κB) is a key transcription factor that is implicated in inflammatory signaling cascade. It plays a significant role in many physiological and pathological processes including inflammation, immunity and metabolism ^19^. Under non-stimulated conditions, NF-κB is inactive in the cytosol as it binds to inhibitor of IκBα. When stimulated using TNFα or with any proinflammatory cytokine, it causes phosphorylation and degradation of IκBα. This releases active NF-κB and causes it to translocate to nucleus followed by the induction of expression of downstream targets ^20^. TNFα is the most classical model to study NF-κB pathway. This TNFα-induced NF-κB activation must be tightly controlled to avoid inflammatory diseases, autoimmunity and cancers ^21,22^. Zhang *et al* have shown that ANGPTL8 acts as a co-receptor of p62 in selective autophagy and that the interaction between p62 and ANGPTL8 are mutually required for IKKγ degradation ^18^. Another group suggested that ANGPTL8 promoted ECM degradation and inflammatory cytokine release through activating NF-κB signaling pathway which displays the detrimental role of ANGPTL8 in intervertebral disc degeneration (IDD). They showed that there was enhanced NF−κB signaling pathway in TNF−α treated Nucleus pulposus (NP) cells through increased expression of ANGPTL8 resulting in ECM degradation and inflammation whereas inhibition of ANGPTL8 resulted in attenuated activation of NF−κB signaling ^23^. Although the role of ANGPTL8 was established in the context of inflammation, the impact of R59W ANGPTL8 variant in regulating inflammation *in vitro* has not been explored so far to our knowledge. Therefore, our objective was to investigate the differential effect of and ANGPTL8 R59W variant over the wild type (WT) in modulating inflammatory pathways.

In this study, we aimed to assess the association of the R59W variant with various clinical lipid and inflammatory traits in a discovery cohort of 867 Arab individuals and a replication cohort of 278 Arab individuals; decipher the functional consequence of these association signals using hepatocyte cell line model under non-stimulated and stimulated TNFα treatment. The impact of overexpressing the ANGPTL8 variant on NF-κB pathway activity and transcriptional activation of the downstream target genes are also assessed. Moreover, structural, and binding dynamics of ANGPTL8 R59W variant and wild type are investigated.

## 2 Materials and Methods

### 2.1 Recruitment of Participants and Study Cohort

The study protocol was reviewed and approved by the Ethical Review Committee of Dasman Diabetes Institute as per the guidelines of the Declaration of Helsinki and of the US Federal Policy for the Protection of Human Subjects. A discovery cohort of 867 participants and an independent replication cohort of 278 participants were recruited. The discovery cohort comprised a random representative adult (>18 years of age) of Arab ethnicity across the six governorates of the State of Kuwait recruited by way of using a stratified random sampling technique for the selection of participants from the computerized register of Kuwait’s Public Authority of Civil Information (PACI). The replication cohort was created by way of recruiting general public that visit our institute for facilities such as physical fitness center and tertiary medical care clinics.

Briefly, native adult Kuwaiti individuals of Arab ethnicity were recruited as study subjects. Pregnant women were excluded. Data on age, sex, health disorders (e.g. diabetes and hypertension) and baseline characteristics such as height, weight, waist circumference and blood pressure were recorded for each participant upon recruitment. Information on whether the participant undergoes medication for lowering lipid levels or for treating diabetes and hypertension was recorded and was subsequently used to adjust the models for genotype-trait association tests. Every participant signed the informed consent form before participating in the study.

### 2.2 Blood sample Collection and Processing

After confirming that the participant had fasted overnight, blood samples were collected in EDTA-treated tubes. DNA was extracted using Gentra Puregene^®^ kit (Qiagen, Valencia, CA, USA) and was quantified using Quant-iT™ PicoGreen^®^ dsDNA Assay Kit (Life Technologies, Grand Island,N NY, USA) and Epoch Microplate Spectrophotometer (BioTek Instruments). Absorbance values at 260–280 nm were checked for adherence to an optical density range of 1.8–2.1.

### 2.3 Estimation of Plasma Levels of Various Biomarkers

Plasma was separated from blood samples by centrifugation, was aliquoted and stored at −80°C. ANGPTL8 was measured as previously reported, ^24,25^, using ELISA kit from (EIAab Sciences, Wuhan, China, Cat# E1164H). Briefly, the samples were diluted ten times using the sample diluent provided in the kit. The standards (recombinant protein of known concentration) provided in the kit was reconstituted and diluted as per kit protocol. Diluted samples and standard were loaded onto the plate and the procedure was followed as per the instructions provided in the kit. The absorbance was read at 450nm on the Synergy H1 plate reader (Bio Tek, Vermont USA) . The concentrations of the unknown samples were determined by Gen 5 (v 5.1) based on a 5 PTL standard curve. TNFα, interleukins and ghrelin were measured using Biolpex multiplex kits (cat# M50-0KCAFOY and cat # 171-A4S01M, Bio-Rad, CA, USA). Standards with known concentrations of specific analytes were reconstituted and diluted as instructed in the manual. The assay was performed, and the plasma levels of the various analytes were determined using the Bio-plex 200 system (Bio-Rad, CA, USA). The concentrations of the unknown samples were calculated by the Bio-plex Manager software (v 6.0) based on a 5 PTL standard curve.

### 2.4 Targeted Genotyping of the ANGPTL8 study variant R59W

Candidate SNP genotyping was performed using TaqMan^®^ Genotyping Assay kit on ABI 7500 Real-Time PCR System from Applied Biosystems (Foster City, CA, USA). Each polymerase chain reaction sample was composed of 10 ng of DNA, 5× FIREPol^®^ Master Mix (Solis BioDyne, Estonia), and 1 μl of 20× TaqMan^®^ SNP Genotyping Assay. Thermal cycling conditions were set at 60°C for 1 min and at 95°C for 15 min followed by 40 cycles of 95°C for 15 seconds and 60°C for 1 min. Sanger sequencing was performed, using the BigDye™ Terminator v3.1 Cycle Sequencing on an Applied Biosystems 3730xl DNA Analyzer, for selected cases of homozygotes and heterozygotes to validate the genotypes determined by the candidate genotyping assay.

### 2.5 Quality check procedures for SNP and Trait Measurements

We used PLINK (version1.9) ^26^ to assess the SNP quality and the trait variance. We calculated minor allele frequency (MAF) and Hardy–Weinberg equilibrium for the study variant. Any quantitative trait value < Q1-1.5 * IQR or any value > Q3+1.5 * IQR was considered as an outlier and was excluded from further statistical analysis.

### 2.6 Allele-based association tests and thresholds for ascertaining statistical significance

Allele-based statistical association tests for the study variant with the quantitative traits and biomarker levels were performed using linear regression analyses adjusting for regular corrections toward age and sex. We also adjusted for diabetes medication and lipid-lowering medication. A threshold of <0.05 was set for P-value.

### 2.7 Cell culture, transfection and treatment

HepG2 cells were procured from American Type Culture Collection (Rockville, Baltimore, MD) and cultured in Minimum Essential Medium (MEM) supplemented with 10% fetal bovine serum (FBS) and penicillin/streptomycin. Cells were seeded to an 80% confluency and proceeded for transfection.

Wild type and R59W clones (Blue Heron Biotech, OriGene, USA) of ANGPTL8 with Myc-DDK tags were used for transfection using Lipofectamine 3000 (Invitrogen, Carlsbad, CA) with 5ug of plasmid for 48hours. Myc tagged pCMV6 vector (OriGene, Rockville, MD, USA) was used as control for the transfection experiments. After 48 hours following transfection, the cells were treated with Recombinant Human TNFα Protein (R&D Systems, MN, USA) at 25ng/mL for 18 hours. The cells were then processed for protein analysis.

### 2.8 Quantitative Real-time PCR

TRIZOL reagent was used for total RNA extraction, and we followed manufacturer instructions to extract RNA. The extracted RNA was then quantified using Epoch spectrophotometer (Biotek, Winooski, VT, USA). The cDNA was prepared from total RNA sample using High-Capacity cDNA Reverse Transcription Kits (Applied Biosystems, Foster City, CA) and it was then amplified with TaqMan gene expression assays or custom primers and normalized to GAPDH using Applied Biosystems 7500 or Rotor-Disc 100 system, respectively. The gene expression analysis was performed using the ΔΔCT method and expressed as fold change. The PCR primers used were TNFα For., 5’-AGGCAGTCAGATCATCTTCTCG-3’; TNFα Rev., 5’-GAGGTACAGGCCCTCTGA TG-3’; IL-10 For., 5’-ACTTTAAGCAAAGGGCTCAGTCC-3’; IL-10 Rev., 5’-GCATCACCTCCTCCAGG TAA-3’;IL-1B For., 5’-AGCAACAAGTGGTGTTCTCCATG-3’; IL-1B Rev., 5’-AAGTGGTAGCAGG AGGCTGA-3’;IL-7 For.,5’-GTCGTCCGCTTCCAATAACC-3’;IL-7Rev., 5’-TCGCGAATTTCCG AATCACC-3’; GAPDH For., 5’-AGGGCTGCTTTTAACTCTGGT-3’; GAPDH Rev., 5’-CCCCACTTGATTT TGGAGGGA -3’. The Human IL-6 and GAPDH primers were taken from TaqMan gene expression assays with ID-Hs00985639_m1 and Hs02786624_g1, respectively.

### 2.9 Western blot analysis

Protein extraction was done using RIPA buffer (50 mM Tris-HCl, pH 7.5, 150mM NaCl, 1% Triton X-100, 1 mM EDTA, 0.5% sodium deoxycholate, and 0.1% SDS) for the cell extracts of transfected cell lysates with/without human TNFα stimulation. Bradford method was used to estimate protein concentrations normalized to β-globulin. 20μg of protein samples were prepared in loading buffer containing β-mercaptoethanol and resolved on 10% SDS-PAGE gels. Then, proteins were transferred onto PVDF membranes (100V for 75 min) and blocked for 2 h at room temperature (RT) using 5% non-fat dried milk in Tris-buffered saline containing 0.05% Tween 20. Membranes were then probed with the primary antibodies P-NF-κB-P65 (3033, Cell signaling), Total NF-κB (8242s, Cell signaling), Phospho-IKKα/β (Ser176/180) (2697, Cell signaling), Total IKKα (2682, Cell signaling), Phospho-IκBα (Ser32/36) (9246, Cell signaling), Total IκBα (9242, Cell signaling) at 4°C for overnight incubation. The membranes were then washed followed by incubation with Rabbit horseradish peroxidase (HRP) conjugated secondary antibody (1:10,000 dilution) for 2 h at room temperature. Protein bands were visualized by chemiluminescence using super sensitivity West Femto ECL reagent (Thermo Scientific, USA), gel images were captured using the Versadoc 5000 system (Bio-Rad, USA), and band intensities were measured using Quantity One Software (Bio-Rad, USA). GAPDH was used as internal control for protein loading and detected using anti-GAPDH antibody (ab2302; Millipore, Temecula, CA, USA).

### 2.10 Luciferase Activity

HepG2 cells were cultured in Minimum Essential Medium (MEM) supplemented with 10% fetal bovine serum (FBS) and penicillin/streptomycin and seeded in 24 well plates. Reporter plasmid carrying luciferase gene under the control of NF-κB response elements was co-transfected with the expression vector encoding ANGPTL8 (pCMV-ANGPTL8), its mutant (ANGPTL8-R59W) and the pCMV empty vector (OriGene Technologies, Inc, Rockville, MD). Luciferase assay (Promega Dual Luciferase assay) was performed 24 hours post transfection as per the instructions in the kit. Briefly, the cells were lysed using the lysis buffer provided in the kit. The lysates were centrifuged for 30 seconds in a refrigerated microcentrifuge. Cleared lysates were used to perform the dual luciferase reporter assay protocol. The cell lysates were mixed with Luciferase reagent II (supplied in the kit) and was measured on Synergy Hybrid H4 plate reader (BioTek, Winooski, VT) and normalized according to protein concentration. Transfection efficiency was assessed using anti-FLAG antibodies as well as against the Renilla luciferase activity.

### 2.11 Structural analysis of ANGPTL8 and binding to IKKβ

The structure of ANGPTL8 was modelled with SWISS-Model ^27^ and I-TASSER ^28^. In this study, we utilized the X-ray crystal structure of iSH2 domain of Phosphatidylinositol 3-kinase (PI3K) p85β subunit (PDB: 3MTT), previously used by Siddiqa *et al*. (^29^ To predict the effect of the R59W substitution on ANGPTL8 structure, DynaMut web, a server that assesses the impact of variants on protein dynamics and stability (Rodrigues, Pires et al. 2021), was used. Furthermore, the modelled ANGPLT8 structure was minimized with YASARA^30^, which finds the lowest energy conformation by reducing the steric energies between bond lengths and angles of both ANGPTL8WT and R59W structures.

The effect of ANGPTL8 variant on the binding to IKKβ was elucidated by interacting both ANGPTL8 WT (R59) and mutant (W59) with IKKβ Prior to modelling the ANGPTL8 protein-protein interactions with IKKβ, the IKKβ interface binding residues for interacting with ANGPTL8 were projected with CPORT ^31^. Protein-protein docking of ANGPTL8 WT (R59) and mutant (W59) with IKKβ (PDB ID: 4KIK) was conducted using the HDOCK server ^32^, which is based on a hybrid algorithm of template-based modelling and ab initio free docking. Model 1, with the lowest docking energy score and the highest ligand RMSD, was selected to analyze binding energy scores (Kd) using the PRODIGY server ^33^. Furthermore, PDBsum ^34^ protein-protein analysis was used to depict interface residues on the secondary structural elements of ANGPTL8 and IKKβ.

### 2.12 Power calculation

We used the Quanto software tool (University of Southern California, Los Angeles, CA, United States) to calculate the power of the study cohorts and its ability to delineate quantitative trait variability at a given power (which was set at 80%). We considered additive mode as the underlying genetic model with “gene only” hypothesis at type 1 error, p ≤ 0.05. Genetic effect that accounts for at least 0.1%– 5% variance in the trait was detected by assuming RG 2 (estimate for marginal genetic effect) values in the range of 0.001–0.05 in step of 0.003. We considered the (mean ± standard deviation) of the quantitative trait and MAF from the study cohorts in these calculations.

## 3. Results

### 3.1 Study cohorts

Two independent cohorts of Arab individuals from Kuwait were recruited for discovery and replication phase. 867 participants formed the discovery cohort and 278 participants formed replication cohort.

### 3.2 Characteristics of the R59W variant

The rs2278426 is a missense (R59W) variant in *ANGPTL8* (betatrophin), while it is an intronic variant in *DOCK6*. The variant downregulates a novel transcript CTC-510F12.4 in whole blood, upregulates *DOCK6* in Adipose-subcutaneous, and is reported to be associated with HDL, LDL, total cholesterol, TG and waist-hip ratio in the NHGRI-EBI GWAS Catalog ^35^. In our study cohorts, the minor allele (T) occurred at a frequency of around 0.010.

### 3.3 Characteristics of the study cohorts

Summary statistics on the clinical characteristics of the participants from the study cohorts are presented in Supplementary Table S1. The mean age of the participants was 43.36±10.78 years in discovery cohort and 46.25±12.38 years in replication cohort. The cohorts comprised mostly class I obese subjects with a mean BMI of 28.53±4.9 (discovery cohort) and 29.93 ±5.17 (replication cohort) kg/m2 and a mean WC of 93.78±11.51 (discovery cohort) and 99.36±13.36 cm (replication cohort). Mean HbA1c (5.47±0.74 and 6.31±1.3% in discovery and replication cohorts, respectively), low-density lipoprotein (3.34±0.89 and 3.13±0.96 mmol/l), high-density lipoprotein (HDL) (1.15±0.29 and 1.2 ± 0.32 mmol/l), total cholesterol (5.19±0.93 and 5.02±1.09 mmol/l) and triglyceride (1.42±0.61 and 1.22± 0.6 mmol/l) were normal or near optimal. Of the subjects from the discovery cohort 38.2% were obese, 29.9% were diabetic, 22.7% were taking medication for diabetes and 22.7% for lowering lipids; these statistics differ considerably with replication cohort -of the subjects from replication cohort 48.7% were obese, 43.3% were diabetic, 36.6% were taking medication for diabetes and 31.8% for lowering lipids.

After partitioning the study cohorts based on the genotypes (CC versus CT+TT) at the rs2278426 variant, the discovery cohort exhibited significant differences (P-value ≤ 0.05) in the mean values for TNFα and IL7 (as well as IL6 and Ghrelin) (Supplementary Table S2); however, the significance of differences in replication cohort are maintained only in the cases of TNFα and IL7 (see Supplementary Table S2 and Supplementary Figure 1).

### 3.4 Association of the *ANGPTL8* rs2278426 variant with increased circulatory levels of TNFα and IL7

The summary statistics of associations observed between the variant and the phenotype traits, derived from allele-based association tests based on additive models corrected for age and sex as well as for medications, are listed in Table 1. The variant was associated with increased levels of TNFα, IL7, IL6 and Ghrelin in the discovery cohort; however, only the associations with TNFα and IL7 were seen replicating. The SNP passed the test for HWE in both the discovery and replication cohorts (Supplementary Table S3).

**Table 1.**
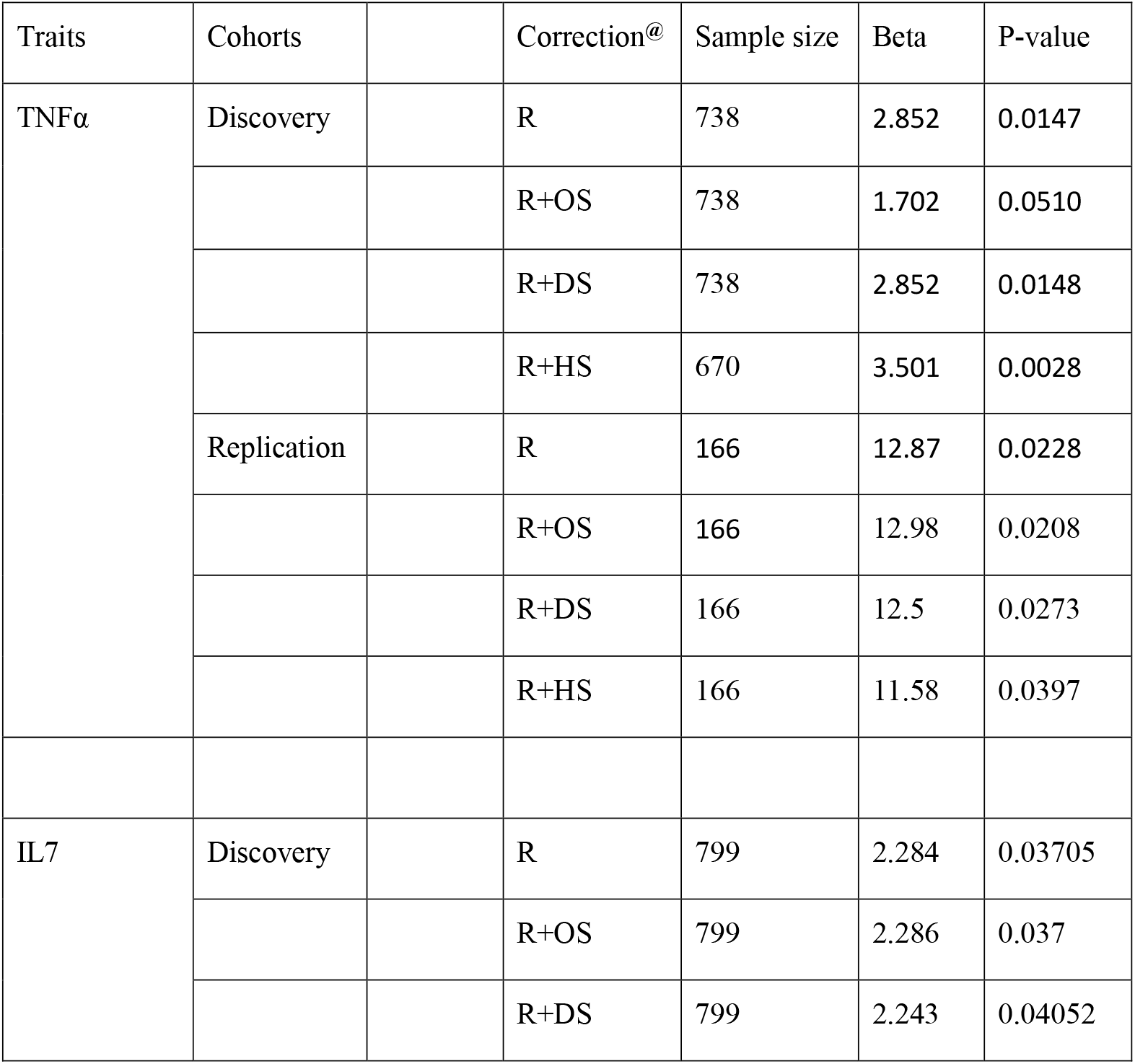
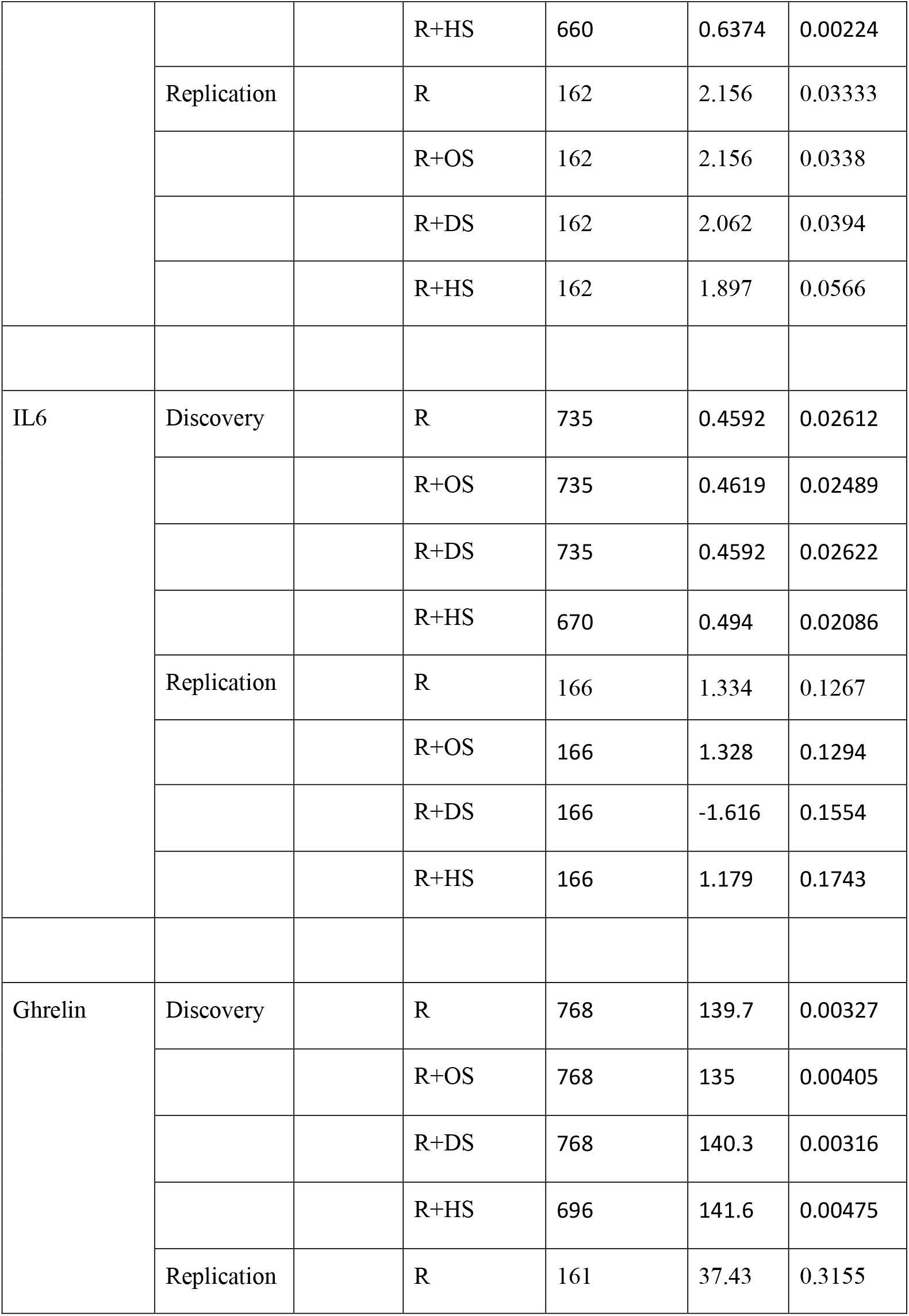
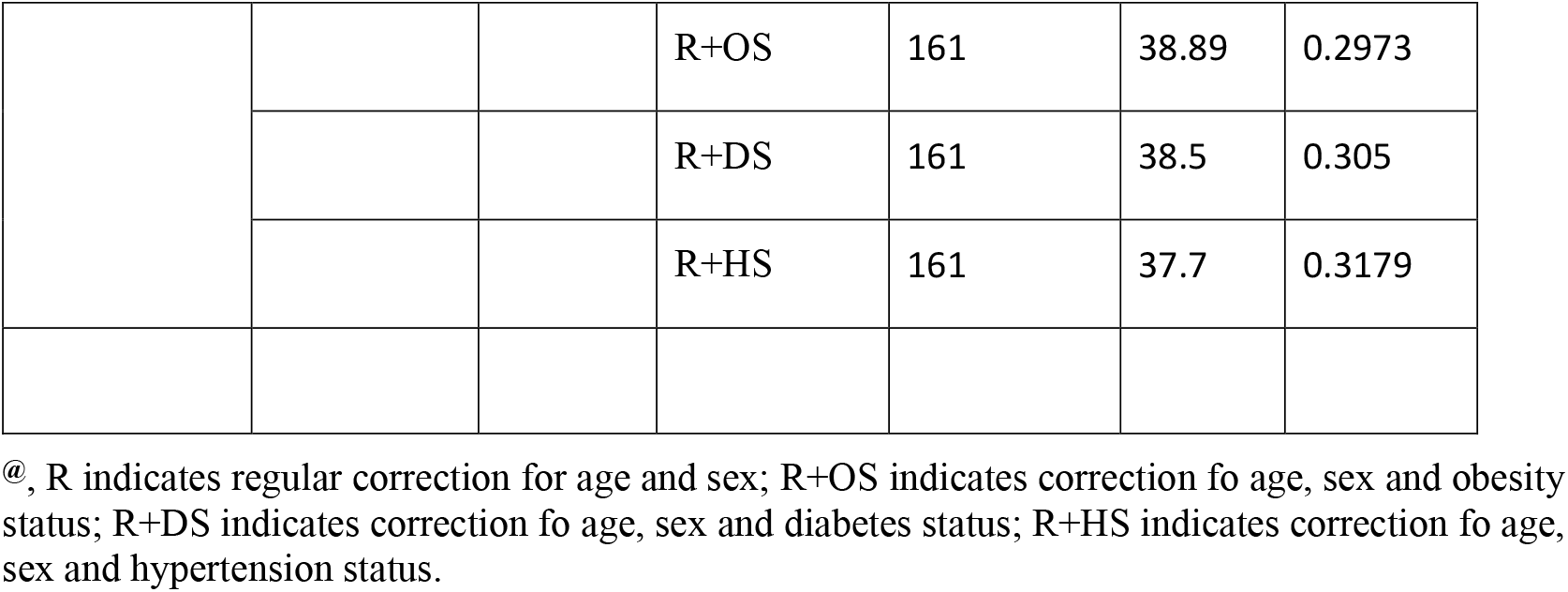
Results of statistical tests for association between the *ANGPTL8* rs2278426 (R59W) variant and biomarkers. Only those associations that showed significant p-values (≤0.05) in the discovery cohort are shown. Of the four biomarkers (TNFα, IL7, Il6 and Ghrelin) that showed significant p-values in discovery cohort, only the TNFα and IL7 showed significant p-values in replication cohort as well.

In order to confirm that the impact by the effect allele at the variant on the levels of TNFα and IL7 is not due to obese or diabetic or hypertensive subjects in the cohort, we performed allele-based logistic regression analysis to evaluate the risk of disorders (diabetes, obesity and hypertension) due to the variant (Supplementary Table S4). Though the odds-ratio (OR) values were notable for the disease status of diabetes and hypertension, the P-values were not significant.

### 3.5 R59W variant activates NF-κB pathway compared to wild type

In order to assess the impact of R59W variant on the activity of NF-κB pathway, we conducted overexpression and luciferase binding assays using HepG2 cell line. A reporter plasmid carrying luciferase gene under the control of NF-κB was co-transfected with the expression vector encoding ANGPTL8 (pCMV-ANGPTL8) or its variant (ANGPTL8-R59W) (OriGene Technologies, Inc, Rockville, MD). Overexpression of the variant resulted in significantly increased phosphorylation of both IKKα/β protein and NF-κB p-65 when compared to the wild type supporting the activation of the NF-κB pathway, p value < 0.05 (Figure 1B, and 1A). Moreover, a significant increase (over 2-folds) in luciferase activity with R59W variant compared to the wild type was observed (Figure 1C). These results indicate that the R59W variant of ANGPTL8 can activate NF-κB pathway compared to the wild type. The expression of the flags for both ANGPTL8 and its variant were comparable following transfection (1D).

**Figure 1.**
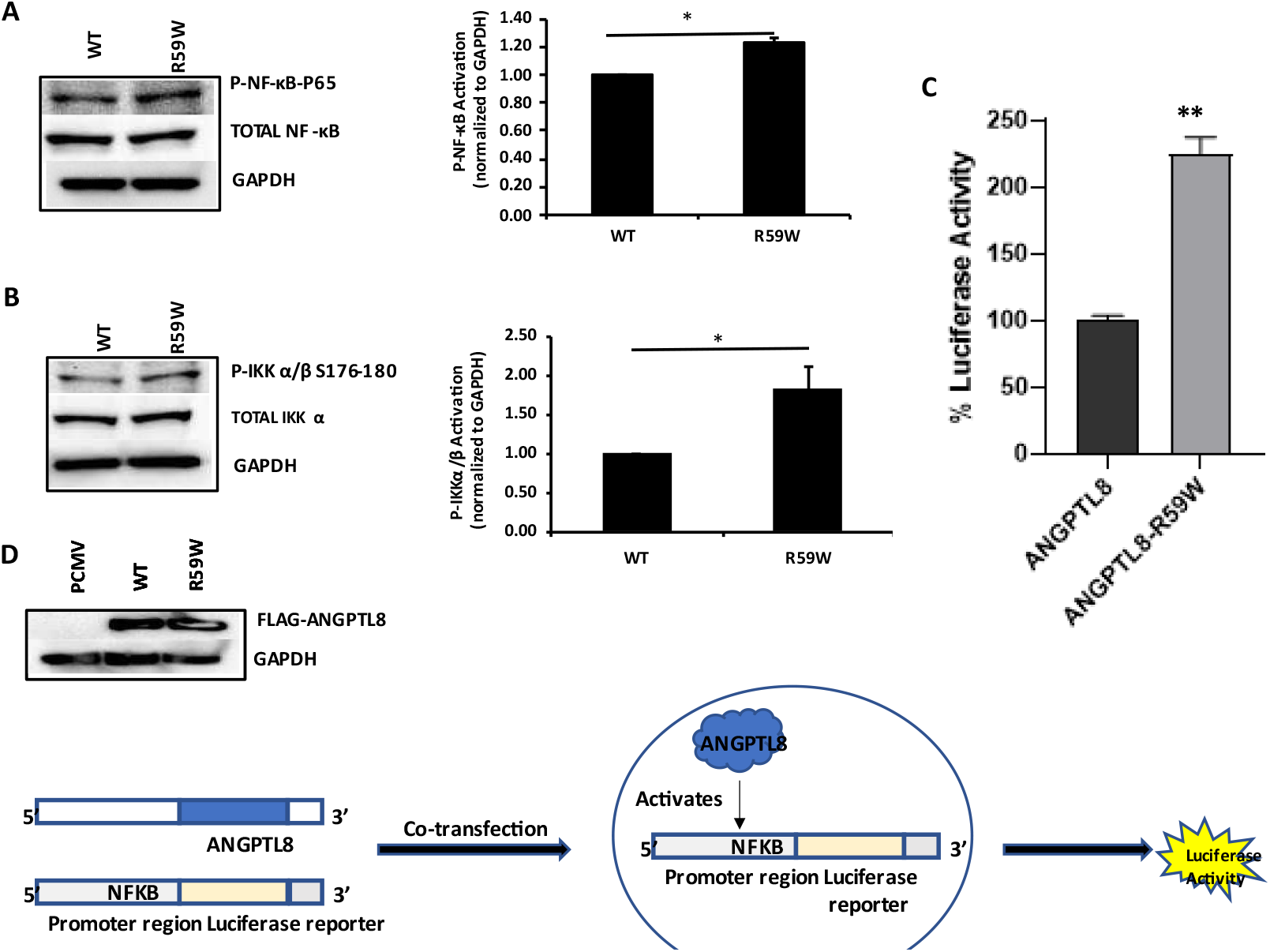
R59W variant activates NF-κB pathway compared to wild type. Overexpression of R59W variant increased the phosphorylation P-NF-KB-P65 (A), P-IKK α/β S176-180 (B), and the activation of NF-κB p65 in the luciferase reporter assay compared to wild type (C), represented image of ANGPTL8 flag and its variant by Western blot (D). *P<0.05,**P<0.001 as determined using student’s t-test, N=4

### 3.6 R59W variant further elevates activation of NF-κB signaling pathway compared to wild type under TNFα stimulation

Literature reports have shown that the transcription and expression of *ANGPTL8* were both significantly increased in HepG2 cells after being treated with TNFα in a dose-dependent fashion ^18^. Therefore, we decided to assess the impact of ANGPTL8 on NF-κB under TNFα stimulation. Under TNFα stimulation, overexpression of the R59W variant resulted in increased phosphorylation of both IKK-α/β protein and NF-κB p-65 compared to the wild type supporting the activation of the NF-κB signaling pathway, P-value <0.05 (Figure 2C,2B, and 2A). In support of this, luciferase reporter assays also revealed significant activation (over 3-folds) of NF-κB p65 by overexpression of R59W variant and subsequent treatment with TNFα (Figure 2D). Altogether, the presented data demonstrated that the R59W variant can potentiate TNFα mediated NF-κB signaling activation compared to wild type.

**Figure 2.**
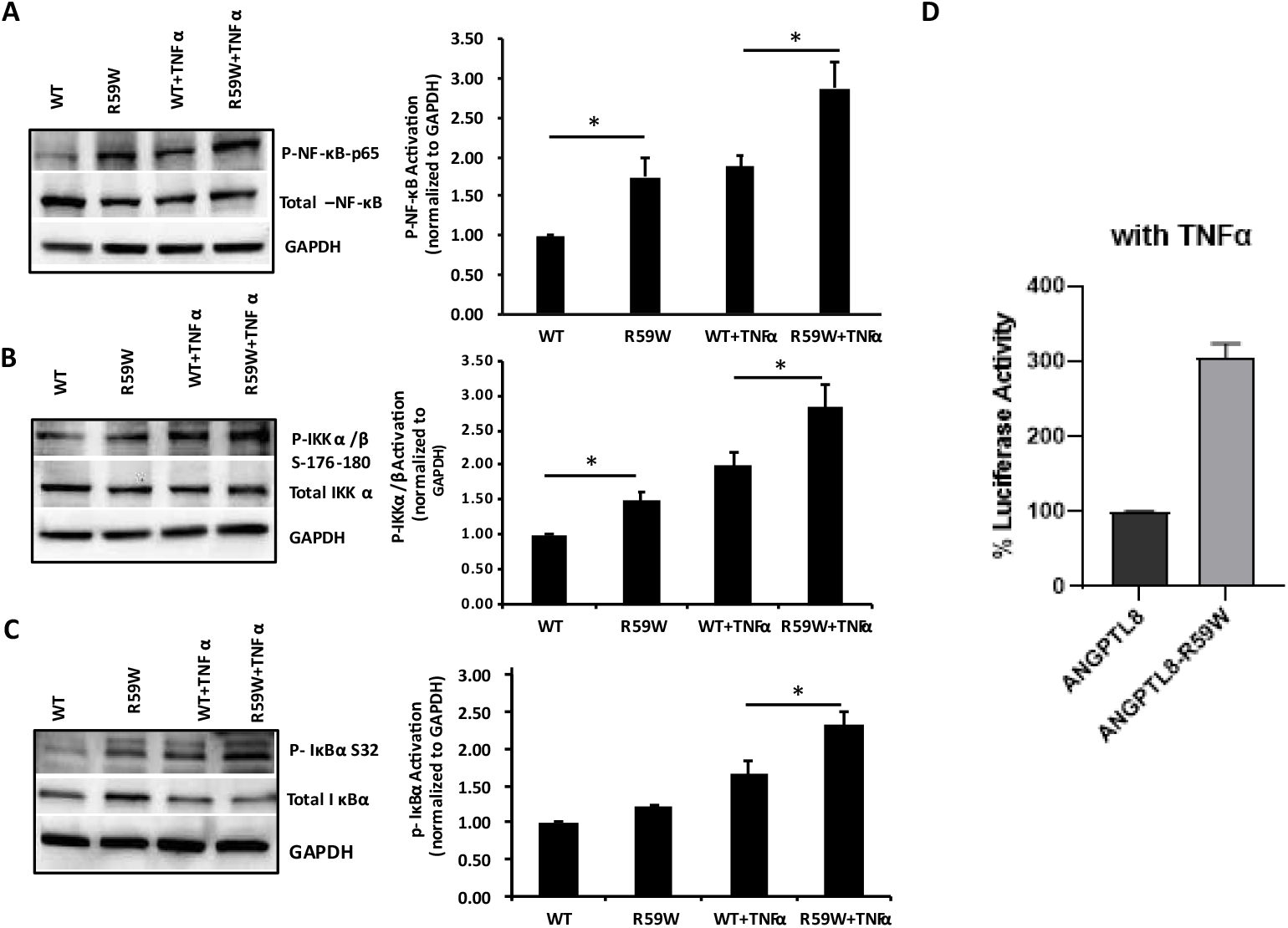
R59W variant activates NF-κB pathway under TNFα stimulation compared to wild type. Overexpression of R59W variant showed increased in the phosphorylation P-IKK α/β S32, S176-180 and P-NF-KB-P65 (A-C), further activation of NF-κB p65 in the luciferase reporter assays when stimulated by TNFα compared to wild type (D) *P<0.05,**P<0.001 as determined using student’s t-test, N=4.

### 3.7 Impact of NF-κB activation on other inflammatory cytokines

To assess and confirm the effect of NF-κB pathway activation, we measured the gene and protein expressions of several inflammatory cytokines (such as TNFα, IL6, IL10, IL-1B and IL7) regulated by NF-κB pathway. There was significant increase in both the gene (Figures 3A, p value <0.05) and protein expression (Supplementary Figure 2) of IL6, TNFα in the R59W variant sample as compared to wild type supporting the impact of NF-κB signaling pathway on the levels of other inflammatory markers.

**Figure 3.**
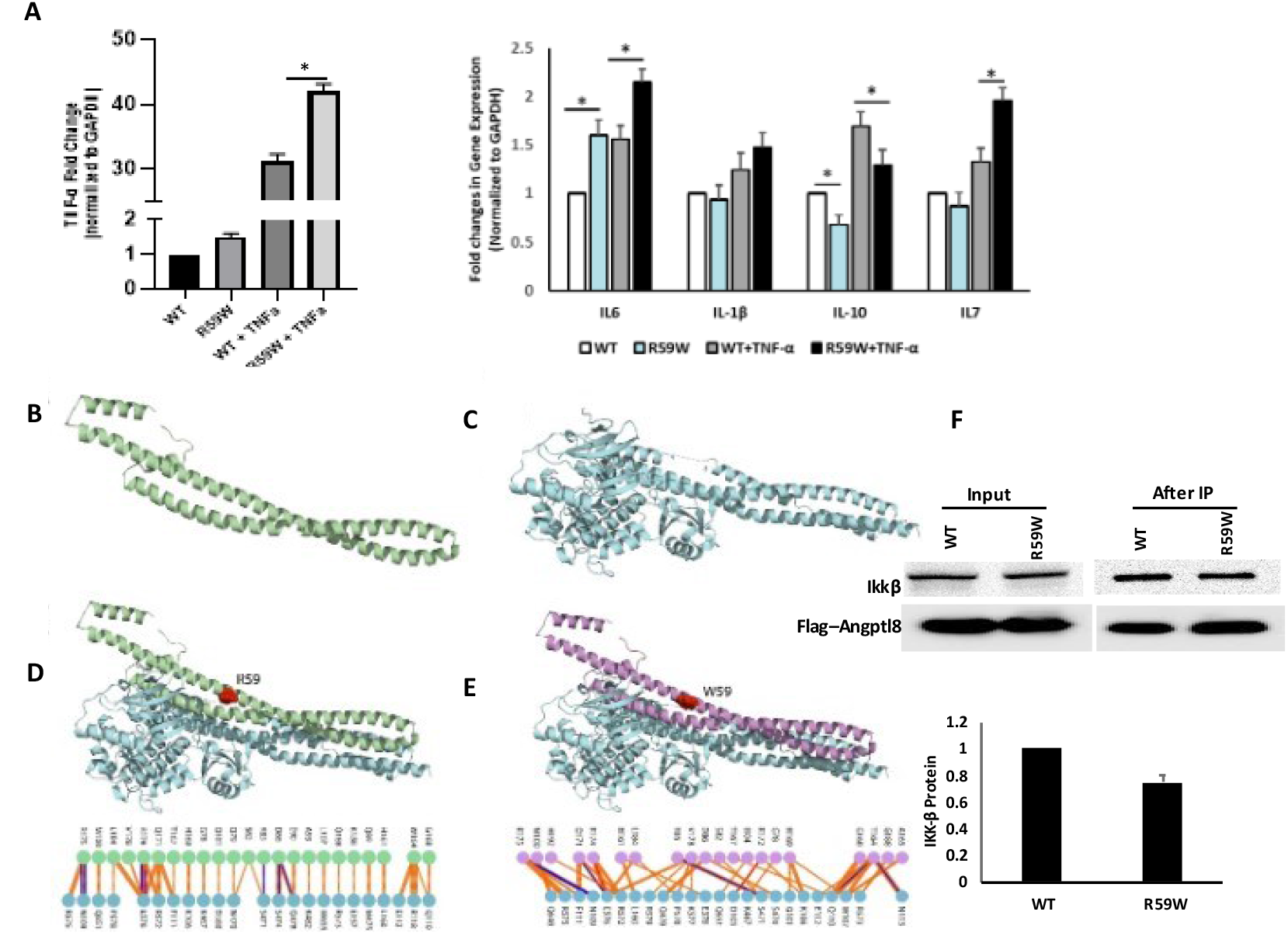
The impact of overexpression of R59W variant on NF-κB downstream target genes and Structural Analysis of ANGPTL8 and IKKβ. Gene expression analyses for several inflammatory cytokines (A), Modelled structure of ANGPTL8 with I-TASSER and Swiss-Model (B) Crystal structure of IKKβ monomer (PDB ID: 4KIK, C). Modelled ANGPTL8 WT (green) with IKKβ (D), Modelled ANGPTL8 W59 (purple) with IKKβ (E), Co-immunoprecipitation assay result on the interaction between ANGPTL8 R59W and IKKβ compared to compared to wild type (F). *P<0.05,**P<0.001 as determined using student’s t-test, N=3.

### 3.8 Structural and binding analyses for ANGPTL8 R59W variant and wild type

The data presented so far demonstrated that ANGPTL8 variant from arginine (R) to tryptophan (W) positioned at residue 59 effect the cellular NF-κB activity. Therefore, understanding the structural changes in the mutated ANGPTL8 is critical to understand its function and potential effects on cellular activity. ANGPTL8 structure modelled with I-TASSER and SWISS-Model demonstrates two α-helical subunits connected by a 6-residue coil (Figure 3B). Furthermore, ANGPTL8 stability analysis using DynaMUT web-server showed that the R59W variant resulted in a more dynamic structure with a ΔΔG of -1.37 kcal/mol.

Zhang *et al*., has suggested that ANGPTL8 binding to IKKβ affects IKKβ cellular function^18^. Therefore, to examine whether the ANGPTL8 R59W variant interaction affected IKKβ downstream cellular function, we modeled both ANGPTL8 WT (R59) and mutant (W59) interaction with IKKβ (Figure 3C) to form an ANGPTL8-IKKβ complex using HDOCK docking server ^32^ (Figures 3D and 3E). The binding affinities of ANGPTL8 R59 and W59 to IKKβ at 37°C were calculated with PRODIGY server. The binding affinity of ANGPTL8 R59 to IKKβ was 4.1 × 10^−8^ Kd/M, whereas ANGPTL8 W59 demonstrated a binding affinity of 3.7 × 10^−8^ Kd/M to IKKβ. Furthermore, the binding interface analysis of ANGPTL8-R59 and -W59 complexes with IKKβ showed different interaction interfaces, whereby ANGPTL8-R59 formed seven H-bonds with IKKβ compared to the six H-bonds formed with ANGPTL8-W59, which further corroborates the weaker binding affinity in the mutant complex with IKKβ in comparison to wild type. Therefore, the weaker interaction demonstrated between the ANGPTL8-W59-IKKβ complex could positively impact NF-κB inflammatory activities presented downstream in comparison to the ANGPTL8-R59 complex with IKKβ (Figure 4). Co-immunoprecipitation assay was performed to support our structural hypothesis on the binding between ANGPTL8 R59W and IKKβ. Weaker binding between ANGPTL8-R59 with IKKβ compared to wild type supporting the structural hypothesis (Figure 3F).

**Figure 4.**
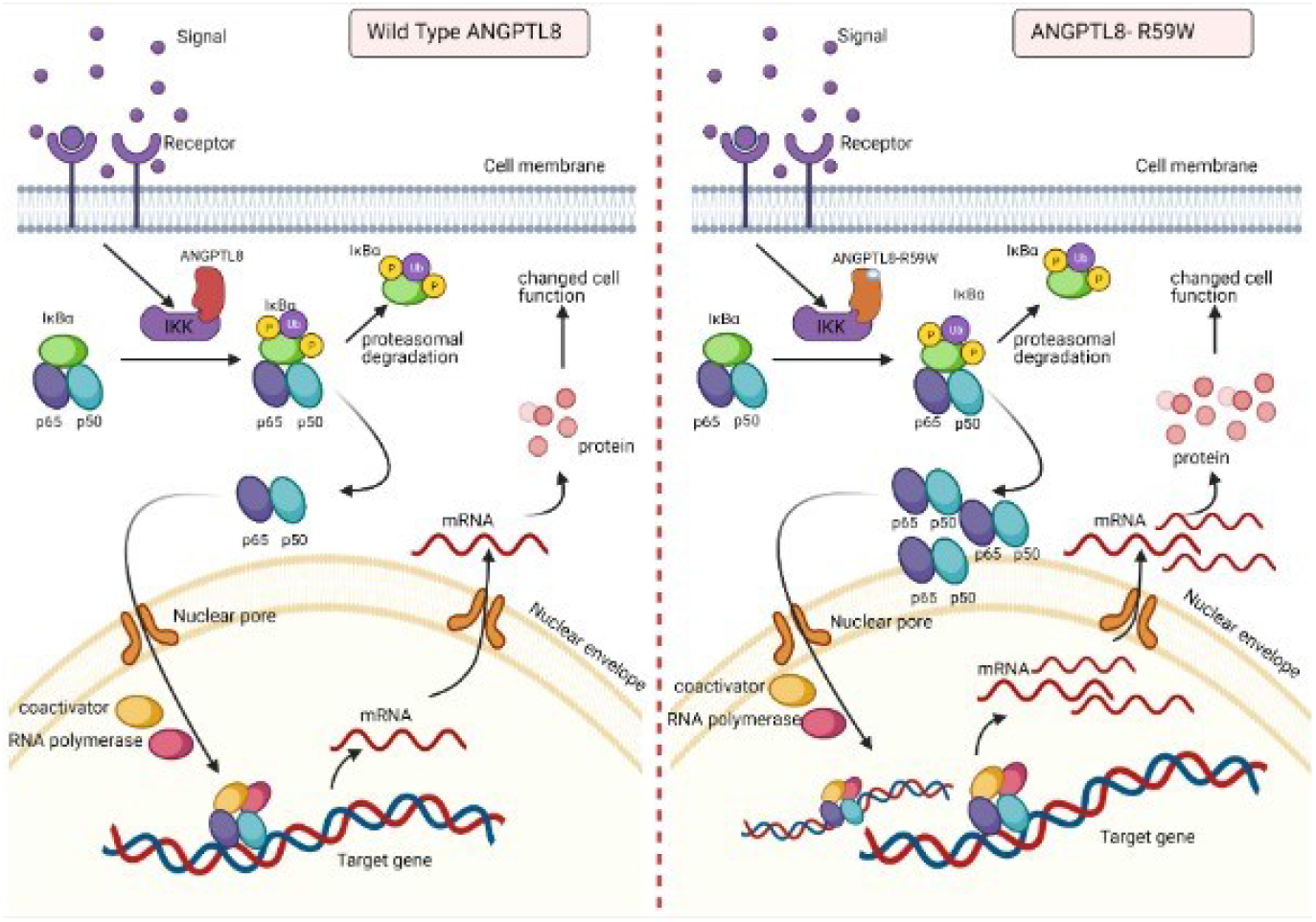
A proposed model for the impact of ANGPTL8 R59W variant overexpression on NF-κB pathway with and without TNFα stimulation.

Further, comparing the binding effect of ANGPTL8 WT (R59) and variant (W59) on the binding to IKKγ (NEMO) was demonstrated using the same modeling techniques explained earlier. Previous studies have shown that the coil region (residues (44-111) is IKKγ binding domain, whereby the coil region was used to model the interaction with ANGPTL8 with the HDOCK docking server ^32^ (Supplementary Figure 3). Moreover, the PRODIGY server predicted the binding affinities of ANGPTL8 R59 and W59 to IKKγ. The binding affinity of ANGPTL8 R59 to IKKγ was 7.8x10-10 Kd/M, whereas ANGPTL8 W59 demonstrated a binding affinity of 3.3x10-9 Kd/M. The weaker binding affinity of IKKγ to ANGPTL8-W59 further indicates that the intermolecular changes in the ANGPTL8 variant affected the dynamics of the protein, which, in turn, reduced the binding affinity to IKKγ. The predicted binding affinities of ANGPTL8 R59 and W59 to IKKγ from the modeled structures are concomitant with the experimental data presented (Supplementary Figure 3).

### 3.9 Results from Power calculation

Results from power calculation indicated that the discovery cohort of the study had 80% power to detect associations with the ANGPTL8 variant explaining 1% variance in the quantitative traits of TNFα of the discovery cohort and 2.8% variance in the replication cohort. The observed effect sizes of TNFα were 2.852 and 12.87 in the discovery and replication cohorts, respectively. These values are closer to the expected effect sizes of 3.271 and 12.628 as revealed by the power calculation for TNFα.

## 4. DISCUSSION

ANGPTL8 plays an important role in lipid and glucose metabolism and is widely associated with various metabolic disorders such as obesity and T2D, with more recent studies highlighting its role in inflammation ^13,36^. However, the role of the *ANGPTL8* rs2278426 (p.R59W) variant remains unclear and needs to be explored in the context of inflammation. To our knowledge, we are the first to investigate the effect of the variant R59W on inflammation. Results from our genetic association tests and *in vitro* assays highlight the influential role of R59W variant of ANGPTL8 in regulating inflammation in hepatocytes. Finally, structural and binding analyses of ANGPTL8 R59W variant revealed a less stable structure with weak transient binding compared to wild type, which possibly explain the increased NF-κB activity in the *in vitro* analyses.

A similar R to W variant has been previously reported and was shown to disrupt functional domains within Troponin T or other proteins ^37^. Our genetic association study has shown that there is an increase in the levels of proinflammatory makers, namely IL-7, IL-6, TNFα and ghrelin, in the blood circulation of individuals having genotypes of alternate allele (CT+TT) as compared to reference homozygotes (CC) in a cohort of 867 individuals. Our previous cross-sectional study of 283 non-diabetic Arab individuals highlighted the significance of R59W variant on glucose metabolism by way of showing that individuals having the R59W variant was associated with higher FBG levels compared to the wild type ^38^. Previous studies have shown that circulating levels of ANGPTL8 was elevated in patients with severe infections and illustrated a strong correlation between ANGPTL8 and LPS-induced acute inflammatory response in animal models ^12^. Circulating levels of ANGPTL8 was higher in human cohorts with NAFLD and it was further elucidated in *in vitro* and *in vivo* models ^17^.

The role of ANGPTL8 in lipid metabolism has been well established. Studies showed that that this genetic variant in *ANGPTL8* is associated with reductions in plasma levels of high-density lipoprotein-cholesterol (HDL-C) in European Americans, Hispanics and African Americans, thus highlighting its role in lipoprotein/TG metabolism ^37^ . Likewise other studies also confirmed the same R59W variant to contribute to lower HDL cholesterol levels in American Indians and Mexican Americans due to increased activation of ANGPTL3 by ANGPTL8 ^39^. Another study of Arab cohort suggested that the gene variants rs737337 (T/C) and rs2278426 (C/T) are associated with lower risk of hypercholesterolemia and hyperglycemia supporting the role of ANGPTL8 in lipid and glucose metabolism ^40^. Additionally, it was shown in Japanese cohorts that the rates of T2D and impaired glucose tolerance were greater in subjects with the R59W variant thereby highlighting its role in diabetes and therefore as a potential target for prevention of T2D ^41^. No group has ever studied the impact of this variant on inflammation.

Emerging evidence suggests that ANGPTL8 may play a role in inflammatory processes. Several studies have investigated the association between ANGPTL8 and inflammation, and while some findings point to a potential involvement, it is important to consider the complexity and limitations of these studies. Further research is required to elucidate the exact mechanisms and the extent of ANGPTL8’s contribution to inflammation in various physiological and pathological contexts. Zhang et al., have shown that ANGPTL8 can work intracellularly as a negative feedback inhibitor of NF-κB activation via self-associating its N-terminal region and interacting with Sequestosome-1 (p62/SQSTM1)^18^. The resulting ANGPTL8-p62 complex aggregates and acts as a platform in the recruitment and selective autophagic degradation of IκB kinase gamma (IKKγ/NEMO)^42^. On the other hand, the results of Liao, Wu *et al*., indicated that ANGPTL8 plays an adverse role in the progression of IDD and that the resulting silencing of ANGPTL8 could reduce ECM degradation and inflammation in human NP cells via inhibition of the NF−κB signaling pathway^23^. These differences were thought to be due to the differences in the cell lines used and the acute nature of stimulation in the first study versus the second. ANGPTL8 could possibly have pleiotropic functions in different tissues and organs; while it acts as a defense mechanism in certain instances, it is rather proinflammatory in chronic inflammation. Our results support the pro-inflammatory role of ANGPTL8 and its variant through modulating NF−κB signaling pathway. Further work is warranted to fully explore the inflammatory role of ANGPTL8 and its variant under different stimulations and in various tissues. Our study further explored the inflammatory pathways linked with the overexpression of wild type and variant (R59W) in hepatocytes with/without TNFα stimulation. Activity levels of NF-κB p65 was higher in the R59W variant as compared to the wild type and it was further elevated with TNFα stimulations for this variant. This data was further supported and reflected by the luciferase activity and confocal microscopy highlighting the impact of R59W in the inflammatory signaling cascade (Figures 2A-E). In the presence of overexpressed R59W variant and under the influence of inflammatory stimuli like TNFα, the activity levels of IKKα/β were enhanced which sequentially phosphorylate the NF-κB inhibitor IκBα and trigger its rapid degradation through the β-TrCP–derived ubiquitin-proteasome pathway, which releases the NF-κB heterodimer from IκBα. No longer repressed by IκBα, the NF-κB heterodimer translocates to the nucleus from the cytoplasm, binds to its downstream target DNA sequence, and induces the expression of many genes involved in immune response (TNFα, IL-1b, IL6, IL10 and IL7) as represented in our hypothetical model (Figure 4). This throws light on the role of R59W in modulating the inflammation through NF-κB signaling pathway due to the exposure of inflammatory stimuli.

Next, we have shown that IL-6 and TNFα expression was consistently upregulated in the R59W overexpressed/TNFα stimulated both at the transcriptional and translational level depicting the proinflammatory role of this variant in hepatocytes. Furthermore, the R59W variant has shown significant changes in modulating differential gene expression of anti/proinflammatory markers such as IL-1B and IL-10. This observation is corroborated by previous studies which have shown that IL-6 can be induced in hepatocytes by LPS treatment through increased translocation of P65 subunit of NF-κB to the nucleus at 1hr of treatment ^43^. A similar study showed that the influence of IL-17 with synergetic role of TNFα contributes to the development of inflammatory liver diseases in primary hepatocytes ^44^. Another study indicated that ANGPTL8 promoted the TNFα-induced ECM degradation and inflammation through the hyperactivation of NF-κB signaling pathway in nucleus pulposus (NP) cells ^23^. Based on this evidence, we speculate that R59W overexpression and stimulation with proinflammatory cytokines like TNFα could potentiate p-IKKα/β activities, inhibiting IκBα and subsequently promoting enhanced nuclear translocation of NF-κB p65 in R59W variant and its downstream targets.

Previous studies have suggested that ANGPTL8 interaction with IKKα/β is necessary for IKKα/β downstream cellular activity. Furthermore, in this study, we demonstrated that the ANGPTL8 W59 variant enhanced NF-κB activity compared to wild type under TNFα stimulation as a result of IKKα/β phosphorylation. Structural modelling of ANGPTL8 wild type and R59W variant presented a helical structure joined by a loop as previously shown by Siddiqa *et al*. ^29^. Stability analysis of ANGPTL8 R59W variant presented a less stable structure than the wild type, whereby such substitutions have been shown to affect protein-protein interaction and lead to differential downstream activities ^45^. Moreover, the effect of the ANGPTL8 W59 variant on the cellular pathway is corroborated by our interaction analysis, which showed that the wild type has a higher binding affinity of 4.1 × 10^−8^ Kd/M to IKKβ compared to ANGPTL8 W59 (3.7 × 10^−8^ Kd/M). This difference in ANGPTL8 WT and W59 binding affinities to IKKβ can influence its cellular activity; it has been previously suggested that ANGPTL8 acts as a transient protein for IKKβ activity ^18^. Proteins involved in transient interactions are essential for many cellular functions ^46^. These interactions are formed and broken easily, and proteins interacting in a weakly transient manner show a fast bound-unbound equilibrium as observed with ANGPTL8-W59-IKKβ complex, resulting in more differential and possibly pronounced downstream activities than ANGPTL8 -WT-IKKβ complex.

To conclude, our studies demonstrate the intracellular and extracellular role of ANGPTL8 R59W variant in regulating proinflammatory events as reflected by genetic association tests, *in vitro* assays, and structural studies. Further studies are warranted to unravel the other signaling pathways and the crosstalk between the ligand-receptor interaction to deepen the knowledge on the mechanistic role of the variant R59W in the context of inflammation. Our current knowledge of the study could provide insight on the role of the R59W variant of ANGPTL8 in inflammation events/cascade. Thus, by modulating the NF-κB signaling pathway through ANGPTL8 R59W variant, it is possible to improve immune response, cell fate and thereby modulate inflammatory diseases.

## 6 Author Contributions

MA: conceptualization, study design, initial manuscript drafting and revision. DM: data analysis and interpretation and initial manuscript drafting and revision. PH: statistical data analysis and manuscript revision. AM: Structural analysis and manuscript revision. AC: data analysis and statistical analysis. SK: Confocal microscopy. NA: Western Blot analysis. FA: data assessment and critically revise the manuscript. IT: data interpretation and critically revised the manuscript. OA: Genotyping analysis. MS: data interpretation and critically revised the manuscript. HA: data interpretation and critically revised the manuscript. RA: data interpretation and critically revised the manuscript. FA: study design, data interpretation, wrote and critically revised the manuscript. TAT: data analysis, statistical analysis, initial manuscript drafting and critical revision of the manuscript. JA: study design, data interpretation, wrote and critically revised the manuscript. All authors have seen and approved the final manuscript.

## 8 Acknowledgments

We are grateful to Clinical Laboratory and the Tissue Bank Core Facility at DDI for their contribution in handling samples. We are also indebted to Kuwait Foundation for the Advancement of Sciences (KFAS) for financial support of this research project (RA AM 2016-003). The corresponding authors had full access to all the data in the study and had final responsibility for the decision to submit for publication.

## 10. Figure Legends

**Supplementary Figure 1. Box plots for TNFa, IL6, and IL7 from the study cohort**.

**Supplementary Figure 2. The impact of overexpression of R59W variant on NF-κB downstream target proteins IL6 and TNFα**(A and B). *P<0.05,**P<0.001 as determined using student’s t-test, N=3.

**Supplementary Figure 3. The 3-dimensional structures of ANGPTL8 WT (red) and R59W (salmon) interacting with IKKγ (A) and Co-immunoprecipitation assay result on the interaction between ANGPTL8 R59W and IKKγ compared to compared to wild type (B)**.

