## Supplement Tables and Figures for "ANGPTL8 R59W variant influences inflammation through modulating NF-κB pathway under TNFα stimulation"

**Supplementary Table S1. Clinical characteristics of the study cohorts**

| Phenotypes | Discovery Cohort | Replication Cohort |
|---|---|---|
| Male:Female | 516:351 (59.5%:40.5%) | 125:153 (45%:55%) |
| Age (years) | 43.36±10.78 | 46.25±12.38 |
| Height (meters) | 1.64±0.86 | 1.64±0.09 |
| Weight (kilograms) | 77.95±15.02 | 81.40±16.23 |
| BMI ($kg/m^2$) | 28.53±4.9 | 29.93±5.17 |
| WC (cm) | 93.78±11.51 | 99.36±13.36 |
| HDL (mmol/l) | 1.15±0.29 | 1.20±0.32 |
| TC (mmol/l) | 5.19±0.93 | 5.02±1.09 |
| LDL (mmol/l) | 3.34±0.89 | 3.13±0.96 |
| TG (mmol/l) | 1.42±0.61 | 1.22±0.59 |
| FPG (mmol/l) | 5.14±0.75 | 5.77±1.24 |
| HbA1c (%) | 5.47±0.74 | 6.31±1.29 |
| Obese:non-obese | 331:536 | 136:142 |
| Diabetics and non-diabetics | 259:608 | 121:157 |
| Hypertensive: non-hypertensive | 364:427 | 84:192 |

| | | |
|---|---|---|
| Diabetes medication | 197:670 | 101:175 |
| Lipid lowering medication | 197:670 | 89:189 |

**Supplementary Table S2. Differences in traits between individuals with reference homozygous genotypes CC and those with carrier genotypes (CT+TT) at the study variant rs2278426 in Study cohort in replication cohort using t-test.**

| **Trait** | **Reference homozygous (CC) in discovery cohort** | **Heterozygous + alternate homozygous (CT+TT) in discovery cohort** | **P-value_ discovery cohort** | **Reference homozygous (CC) in replication cohort** | **Heterozygous + alternate homozygous (CT+TT) in replication cohort** | **P-value_ replication cohort** |
|---|---|---|---|---|---|---|
| TNFα | 33.39±13.52 | 36.47±13.26 | 0.012 | 124.9±31.3 | 138.7±33.1 | 0.03 |
| IL7 | 6.42±2.36 | 6.99±2.39 | 0.011 | 12.7±5.7 | 14.9±5.6 | 0.05 |
| IL6 | 6.60±2.41 | 7.04±2.21 | 0.041 | 16.70±4.61 | 18.06±5.08 | 0.161 |
| IL4 | 2.35±0.77 | 2.39±0.71 | 0.56 | 6.15±1.68 | 6.40±1.85 | 0.478 |
| Ghrelin | 1077.9±547.5 | 1250.6±630.3 | 0.002 | 536.83±208.6 | 575.54±236.2 | 0.401 |

**Supplementary Table S3. SNP quality assessment tests on the ANGPTL8 rs2278426 (R59W) for HWE**

| Cohort | Minor/Major alleles | MAF | Genotype counts | Observed heterozygous | Expected heterozygous | P-value HWE |
|---|---|---|---|---|---|---|
| Discovery | T/C | 0.102 | 11/155/701 | 0.178 | 0.183 | 0.457 |
| Replication | T/C | 0.101 | 5/46/227 | 0.166 | 0.181 | 0.173 |

**Supplementary Table S4: Logistic regression analysis for impact of the variant rs2278426_T on the disease status of the participants in the cohorts**

| Disease status | Cohorts | OR [CI] | Standard error | P-value |
|---|---|---|---|---|
| Diabetes status | Discovery cohort | 1.15 [0.80-1.64] | 0.181 | 0.442 |
| | Replication cohort | 1.22 [0.68-2.19] | 0.296 | 0.486 |
| Obese status | Discovery cohort | 0.898 [0.64-1.24] | 0.166 | 0.521 |
| | Replication cohort | 0.93 [0.54-1.61] | 0.274 | 0.812 |
| Hypertension status | Discovery cohort | 1.06 [0.76-1.49] | 0.171 | 0.709 |
| | Replication cohort | 1.27 [0.68-2.35] | 0.315 | 0.447 |

**Supplementary Figure 1. Box plots for TNFa, IL6, and IL7 from the study cohort.**

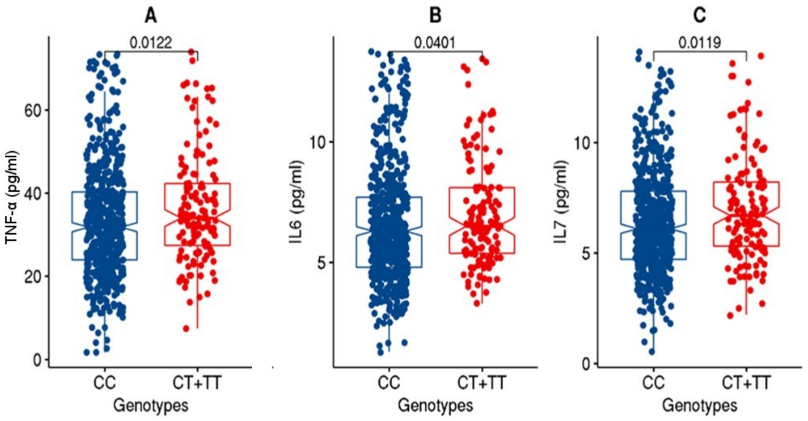


**Supplementary Figure 1**

**Supplementary Figure 2. The impact of overexpression of R59W variant on NF-κB downstream target proteins IL6 and TNFα** (A and B). *P<0.05,**P<0.001 as determined using student's t-test, N=3.

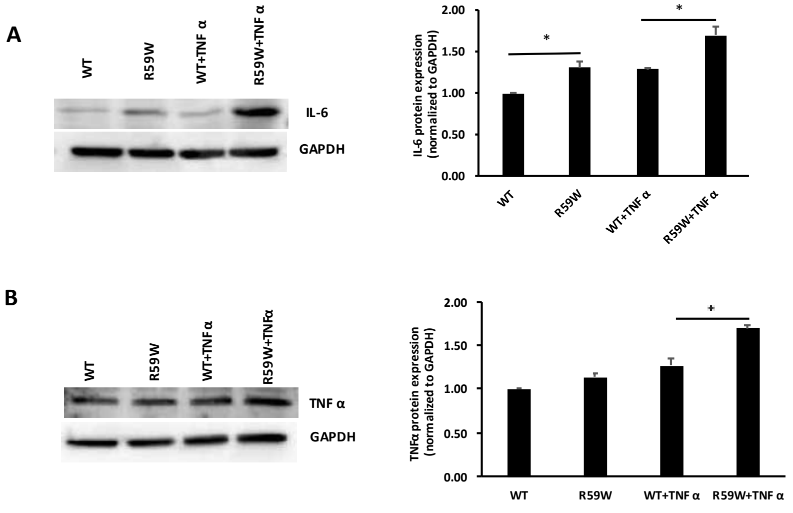


**Supplementary Figure 2**

**Supplementary Figure 3. The 3-dimensional structures of ANGPTL8 WT (red) and R59W (salmon) interacting with IKKɣ (A) and Co-immunoprecipitation assay result on the interaction between ANGPTL8 R59W and IKKɣ compared to compared to wild type (B).**

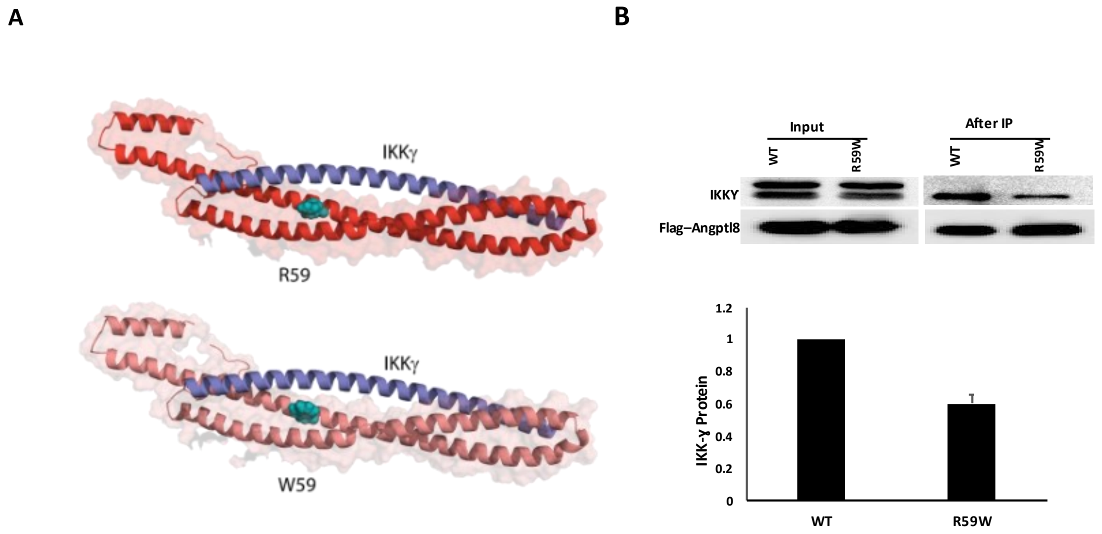


**Supplementary Figure 3**
